# Evolution of threat response-related polymorphisms at the *SLC6A4* locus in callitrichid primates

**DOI:** 10.1101/2023.12.09.570946

**Authors:** Hanlu Twyman, India Heywood, Marília Barros, Jorge Zeredo, Nicholas I. Mundy, Andrea Santangelo

**Author notes:** Address for correspondence: Dr. Nicholas I. Mundy, Department of Zoology, University of Cambridge, Cambridge, CB2 3EJ. Joint senior authors (Nicholas I. Mundy, Andrea Santangelo).

## Abstract

Variation in an upstream repetitive region at the *SLC6A4* locus, which encodes a serotonin transporter, is associated with anxiety-related behaviour in a few primate species, including humans and rhesus macaques. In this study we investigate evolution of *SLC6A4* polymorphisms associated with anxiety-related behaviour in common marmosets (*Callithrix jacchus*). Assaying variation in the *SLC6A4* repeat region across 14 species in 8 genera of callitrichid primates (marmosets and tamarins) we find large interspecific variation in the number of repeats present (24-43). The black tufted-ear marmoset (*C. penicillata*) has sequence polymorphisms similar to those found in the common marmoset, which is its sister species, and no other species has intraspecific variation at these sites. We conclude that, similar to humans and rhesus macaques, the functional polymorphism at *SLC6A4* in common marmosets has a recent evolutionary origin, and that the anxiety-related allele is evolutionarily derived. Common/black tufted-ear marmosets and rhesus macaques share high ecological adaptability and behavioural flexibility that we propose may be related to the maintenance of the polymorphism.

## Introduction

Interest in the genetic architecture of behaviour and its evolution has increased in recent years. Behavioural variation is generally expected to be highly polygenic (McKay 2009), and this is considered particularly likely for complex behaviours in organisms with a highly developed behavioural repertoire. However, in some cases behavioural polymorphisms occur, where genes of major effect control a large proportion of behavioural variance (e.g. de Belle et al. 1989, Krieger & Ross 2002). These cases are of particular interest since they raise questions about how such polymorphisms could be adaptive, and how they are maintained, in some cases over long spans of evolutionary time.

Behavioural polymorphisms appear to be relatively rare in terrestrial vertebrates, although there are some prominent cases involving mating systems in birds (Thomas et al. 2008, Küpper et al. 2016, Lamichanney et al. 2016). Among mammals, a striking example of a genetic polymorphism affecting behaviour occurs at the *SLC6A4* locus, which encodes a serotonin (5-HT) transporter that plays a critical role at serotonergic synapses in the brain. In three unrelated primate species – humans, rhesus macaques (*Macaca mulatta*) and common marmosets (*Callithrix jacchus*) – polymorphism in a repeat region upstream of the *SLC6A4* coding region is associated with anxiety and emotional reaction to threat stimuli (Lesch et al. 1996, 1997, Santangelo et al. 2016). In all cases, alleles associated with enhanced fear-related behaviour showed a reduced expression of *SLC6A4*. The variants linked with behaviour vary among the species. In humans, functional variants comprise length variants caused by variable numbers of repeats (“short” and “long” alleles), and a SNP in the long allele (Lesch et al. 1996, Hu et al. 2006). In rhesus macaques, there are short and long alleles (Lesch et al. 1997). In common marmosets, the length of the repeat region is invariant, and there are multiple SNPs in the repeat region associated with behaviour (Santangelo et al. 2016).

Key questions concerning these polymorphisms are their evolutionary origin and nature of selection acting on them. Comparative studies have shown that the functional polymorphisms have recent evolutionary origins in humans and rhesus macaques. In great apes, there is variation in the length of the repeat region among species but no evidence for intraspecific polymorphism in repeat length as occurs in humans, implying that the functional polymorphism evolved uniquely along the human lineage (Inoue-Murayama et al. 2000, Claw et al. 2010). Similarly, the polymorphism found in rhesus macaques is absent from other macaques (Chakraborty et al. 2010, Shattuck et al. 2014, Kalbitzer et al. 2016). Furthermore, in a careful study of long-tailed macaques (*M. fascicularis*), which are closely related to rhesus macaques, no associations were found between behaviour and genetic variation at any sites in *SLC6A4* (Miller-Butterworth et al. 2007). The restricted presence of a functional *SLC6A4* polymorphism in macaques has led to speculation about the selective pressures that may be unique to rhesus macaques, with discussion focusing around levels of aggression and ecological success (Wendland et al. 2006, Chakraborty et al. 2010).

The association between *SLC6A4* polymorphism and anxiety-related behaviour in common marmosets was described recently (Santangelo et al. 2016, 2019). In common marmosets, the *SLC6A4* repeat region contains four SNPs that define two major haplotypes that are strongly associated with measures of the emotional response of adult monkeys to a human intruder (Santangelo et al. 2016). In addition, the polymorphism is associated with differential expression of serotonin 2A receptors and microRNAs in brain areas underlying emotional processing (Santangelo et al. 2019, Popa et al. 2022). This first functional *SLC6A4* polymorphism in a New World primate is fascinating since it may shed light on the adaptive origins of these polymorphisms across primates. The polymorphism has been confirmed in a wild population of common marmosets (Santangelo et al. 2016) but there is currently little knowledge of its evolutionary origin among the New World (platyrrhine) primates.

Marmosets are members of the callitrichid clade of New World monkeys, which comprise nine genera of small monkeys with a broad range of ecological niches and social systems. In particular, callitrichids occupy a range of forest types, from the moist lowland forest of Amazonia, to the Atlantic forest of coastal Brazil, to dry forests of northeastern Brazil, which is one of the native habitats of the common marmoset. The distribution of the *SLC6A4* polymorphism in relation to variation in socioecology of callitrichids could provide important clues to its maintenance.

Here we investigate the evolution of the common marmoset *SLC6A4* polymorphisms by re-sequencing the repeat region across a broad range of callitrichid primates, including all major genera. We document for the first time the structure of the repeat region across callitrichids. We ask when the functional polymorphisms in the common marmoset evolved, and whether the anxiety-related allele was ancestral or derived. We consider how the functional polymorphism is distributed in relation to ecological factors, and consider whether common selective pressures could plausibly be related to the distribution of functional *SLC6A4* polymorphisms in callitrichids and macaques.

## Materials and methods

### Samples

Samples are listed in Supplemental Table 1. We sampled broadly across the callitrichid phylogeny, including eight out of nine currently recognised genera (tufted-ear marmosets - *Callithrix*; pygmy marmosets – *Cebuella*; bare and tassel-ear marmosets – *Mico*; Goeldi’s monkey – *Callimico*; lion tamarins - *Leontopithecus*; tamarins - *Leontocebus, Oedipomidas Tamarinus, Saguinus*; Brcko et al. 2022). We sampled multiple individuals per species where possible in order to increase the possibility of detecting allelic variants.

### Laboratory methods

Genomic DNA was extracted from hair and tissue samples using Qiagen kits according to the manufacturer’s protocol. Primers (Supplemental Table 2) were used to amplify the full length of the upstream repeat region of *SLC6A4* (∼800-1200bp). Amplifications were carried out in 25μl reaction volumes containing 0.125μl HotStarTaq (Qiagen), 1x Q-solution, 200 μM of each dNTP, and 0.4 μM of each primer. After 5 min of initial denaturation at 94 °C, each of the 40 cycles consisted of the following steps: 94 °C for 30 s, 60-65 °C for 45 s, and 72°C for 90 s, followed by a 5-min final extension. Excess primers and nucleotides were either removed using the QIAGEN purification kit or EXOSAP-IT (Amersham Biosciences). All amplification products were Sanger sequenced on both strands (with primers SeqF2, SeqR1, RPRRL and SeqF1) at the Sequencing Facility, Biochemistry Department, University of Cambridge and Sequencing Lab at the Catholic University of Brasilia, Brasilia.

The new sequences reported in this study were deposited in Dryad (URL:) and GenBank (Accession nos: …).

### Data editing and analysis

Sequences were edited in DNASTAR and aligned using Muscle in MEGA-X (Kumar et al. 2018). Reconstruction of *SLC6A4* sequence evolution was performed using parsimony.

## Results

### Structure of the SLC6A4 repeat region in callitrichids

We obtained new data from 67 individuals of 14 species of callitrichids (Suppl Table 1). All species contain a *SLC6A4* repeat region. The number of repeats varied substantially among species (range 24 – 43), and there is much sequence variation among repeats in the same species. Repeat number showed low variation within genera and more variation between genera (*Callithrix* 32; *Cebuella* 30-31; *Mico* 25; *Callimico* 34; *Leontopithecus* 42-43; *Leontocebus* 24-25; *Oedipomidas* 28; *Tamarinus* 29; *Saguinus* 26-27).

### Reconstruction of functional SLC6A4 polymorphisms found in common marmoset

The homologous nucleotide sites for the four function-related SNPs in repeats 3, 4 and 23 in common marmosets, defining the CT/T/C and AC/C/G haplotypes, could be unambiguously identified in most species (Figure 1). In the silvery marmoset (*Mico*) and tamarin genera (*Saguinus, Leontocebus, Oedipomidas, Tamarinus*), the repeat homologous to repeat 4 in other callitrichids appears to be absent (Figure 1).

**Figure 1.**
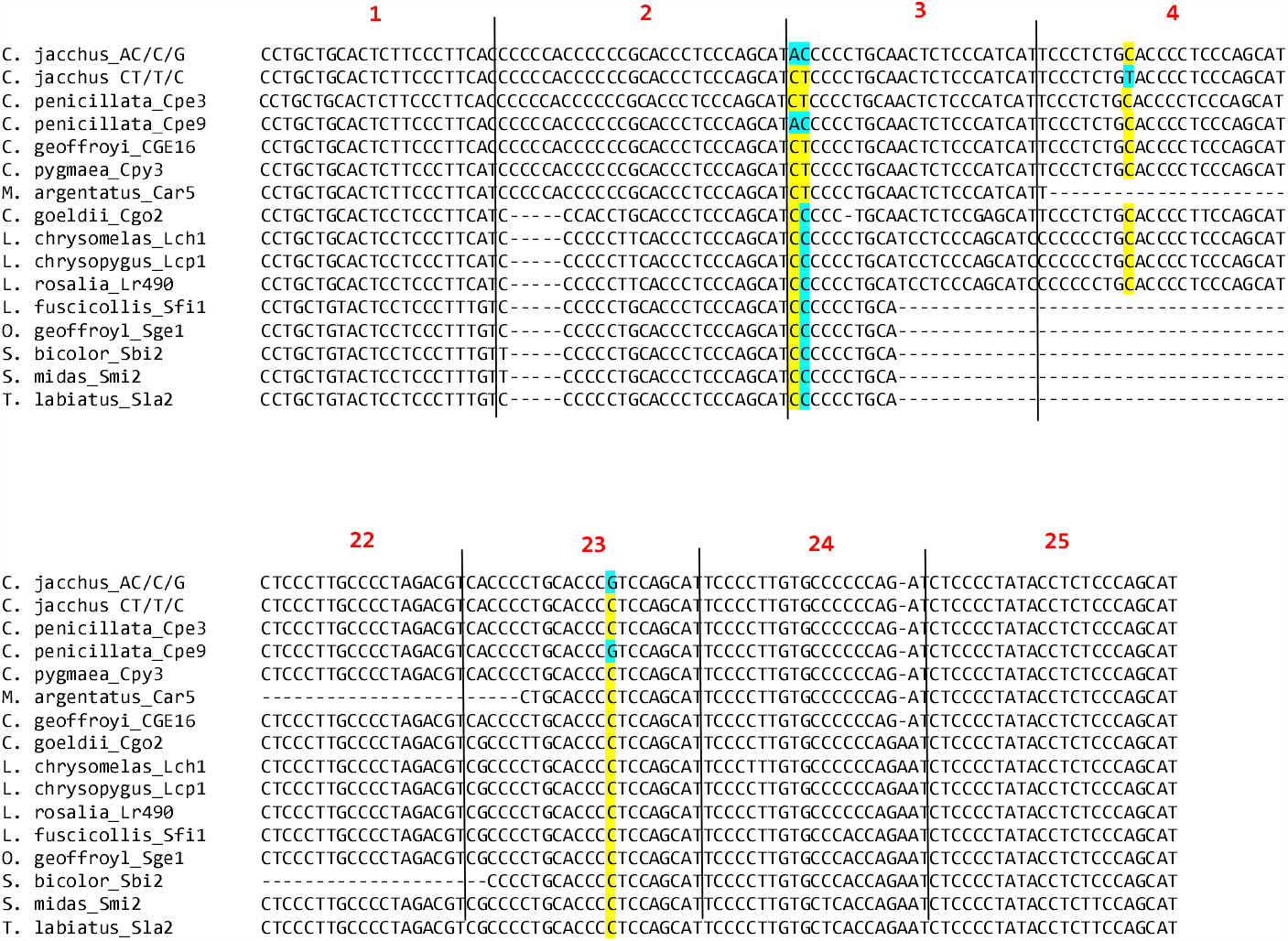
Alignment of common marmoset functional SNPs across species. Alignments of *SLC6A4* sequences encompassing the four SNP sites (highlighted) in the common marmoset that are associated with behaviour, from representative individuals of all species in the study. Repeats are numbered with respect to the common marmoset sequence.

Across the dataset only one species apart from the common marmoset shows evidence of intraspecific polymorphism at any of the four behaviour-related SNPs and this is the black tufted-ear marmoset (*Callithrix penicillata*), the sister species of the common marmoset. Three of the four sites in the black tufted-ear marmoset are variable and these define two haplotypes, CT/C/C and AC/C/G, one of which (AC/C/G) is shared with the common marmoset, and the other (CT/C/C) has a single transition compared to the common marmoset (Figure 1). Notably, none of the four sites are variable in Geoffroy’s marmoset (*C. geoffroyi*, N = 9), which is closely related to the common marmoset and black tufted-ear marmoset, and the single haplotype present in this species is CT/C/C. Among the seven individuals of black tufted-ear sampled, two were homozygous for the AC/C/G haplotype, four were homozygous CT/C/C, and one was heterozygous, giving the following haplotype frequencies: AC/C/G: 0.36; CT/C/C: 0.64.

A reconstruction of the evolutionary history of the four sites is shown in Figure 2. Over most callitrichids these four sites are quite well conserved. The ancestral state for the clade containing marmosets (*Callithrix, Mico, Cebuella*), Goeldi’s monkey (*Callimico*) and lion tamarins (*Leontopithecus*) is reconstructed as CC/C/C, and this haplotype is retained in *Callimico* and *Leontopithecus*. A single transition to CT/C/C occurs in the ancestral marmoset clade (genera *Mico, Cebuella, Callithrix*). Further evolution then occurs in the common ancestor of common and black tufted-ear marmosets, where two derived polymorphisms arise (A/C at the 1st position and C/G at the 4th position), which are the only transversions among the four sites detected in callitrichid evolution. The AC/C/G haplotype, which is associated with anxiety-related behaviour in common marmosets, is clearly the more derived haplotype. Finally, a further transition occurs in one common marmoset haplotype (CT/C/C to CT/T/C).

**Figure 2.**
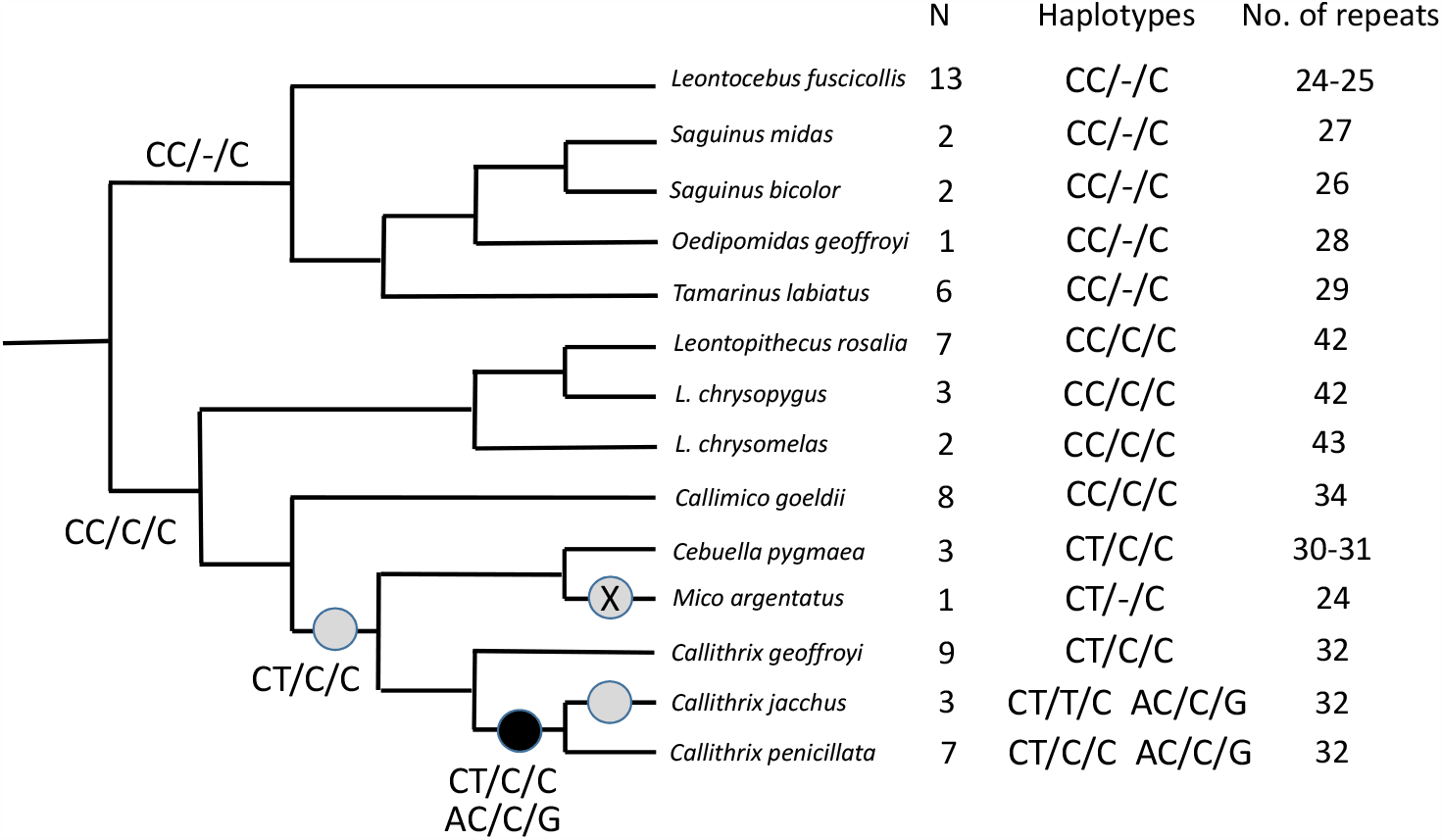
Phylogeny with reconstruction of evolution at the sites of functional SNPs in the common marmoset. Phylogeny of species in the study, with reconstruction for the four SNP sites associated with behaviour in the common marmoset, followed by sample size, haplotypes and number of repeats in repetitive region. The circles indicate reconstructed mutational events – grey = transition, black = origin of functional polymorphism, cross = deletion. The phylogeny is taken from Perelman et al. (2011), with extra information for lion tamarins from Perez-Sweeney et al. (2008) and tufted-ear marmosets from Malukiewicz et al. (2017).

### Intraspecific variation in SLC6A4 in other callitrichids

Variation in other parts of the repeat region of *SLC6A4* was common in other species (Supplementary Table 3). Notable findings were variation in the number of whole length repeats in the pygmy marmoset (*Cebuella pygmaea*) and the saddleback tamarin (*Leontocebus fuscicollis*).

## Discussion

Here we have investigated the evolution of functional behavioural polymorphism at the *SLC26A4* locus in callitrichid primates. Focusing on the SNPs that are associated with anxiety-related behaviour in common marmosets, we find that the black-eared marmoset, the sister species of the common marmoset, is the only other polymorphic species. The anxiety-related haplotype, AC/C/G, is present in these two species and is clearly evolutionarily derived. The *SLC26A4* locus repeat region structure is divergent among callitrichid genera, and intraspecific variation in the repeat region is common. We now discuss the implications of these findings for the evolution of behavioural polymorphism at this locus.

The pattern of variation in *SLC26A4* is similar in the black tufted-ear and common marmosets, with three out of four differences between the two major haplotypes shared. We conclude that the functional polymorphism pre-dated the split between these species and this strongly suggests that it is maintained by some form of balancing selection. This is supported by the substantial frequencies of the two haplotypes in the two species - 0.34 AC/C/G, 0.66 CT/T/C in wild common marmosets (Santangelo et al. 2016) and 0.36 AC/C/G, 0.64 CT/C/C in black tufted-ear marmoset (this study). It will be important to confirm whether variation at these haplotypes is associated with behavioural variation in black tufted-ear marmosets.

In contrast, none of the four sites were variable in any other callitrichid species examined. Of particular interest is the lack of polymorphism in Geoffroy’s marmoset, a close relative of common and black tufted-ear marmosets (Malukiewicz et al 2017). This was based on a relatively small sample size (N = 9), but it is notable that this sample does not appear to have low genetic variation overall since these same individuals are polymorphic for all three alleles at the X-linked opsin locus (see below, Mundy & Caine 2000). If this is a representative sample, there is a >99% chance of detecting the polymorphism with a minor allele frequency of ≥ 0.25, comparable to that in the common and black tufted-ear marmosets. It is therefore likely that the functional polymorphism arose after the split between Geoffroy’s marmoset and the common ancestor of common and black tufted-ear marmosets, although we cannot rule out an earlier origin.

The relatively recent evolution and restricted distribution of functional polymorphic sites at *SLC6A4* in callitrichids is in strong contrast to the pattern seen with another functional polymorphism, at the X-linked opsin locus that underlies individual variation in colour vision. The X-linked opsin polymorphism is pervasive across callitrichids, and, moreover, most species have the same three functional alleles, defined by SNPs in exons 3 and 5 of the opsin coding sequence (Surridge & Mundy 2002). There is widespread trans-specific evolution of X-linked opsin alleles in callitrichids, and presumably similar selective forces maintaining variation in different species. Functional polymorphism at *SLC6A4* shows a different pattern of variation unique to a small clade of marmosets.

Behavioural studies in other species will be needed to determine whether SNPs and/or length variants in other parts of the *SLC6A4* repeat region documented here are functionally relevant. The degree of intergeneric variation in number of repeats is greater than that reported in other primates (e.g. great apes and papionine monkeys; Inoue-Murayama et al. 2000, Kalbitzer et al. 2016), and the arrays of 42-43 repeats found in lion tamarins appear to be the longest recorded. However, the functional consequences of intergeneric variation in the number of repeats are unclear.

Ecologically, it is interesting to note that the two callitrichid species that share the functional *SLC6A4* polymorphism occupy a variety of habitats, some of which are arid, and that they both have relatively large ranges in eastern and north-eastern Brazil, among the largest of any callitrichids. For example, the common marmoset occupies both dry caatinga and moist Atlantic rainforest, and shows considerable behavioural flexibility among these habitats (Abreu et al. 2016, but see Garber et al. 2019). In addition, these species can thrive in modified urban environments (Pontes & Soares 2005, Nogueira dos Santos et al 2014, Secco et al 2018), and they have established permanent populations outside of their normal range in other parts of Brazil (e.g. Cunha et al. 2006). An interesting possibility is that the evolution of the *SLC6A4* polymorphism in these two species of tufted-ear marmosets is related to their ecological flexibility and adaptability.

The overall pattern of functional polymorphic variation in *SLC6A4* uncovered in callitrichids shares several similarities to that in macaques. In both clades a restricted number of species show the polymorphism, the polymorphism likely has a recent origin, and the anxiety-related allele is derived. Rhesus macaques have a large range and a more temperate distribution in general than most other macaque species, and it has been previously suggested that the *SLC6A4* polymorphism may be related to successful exploitation of this habitat. The detailed adaptive mechanisms are poorly understood but a number of studies have shown association of ecologically relevant traits with the polymorphism, e.g. male dispersal (Trefilov et al 2010) and juvenile development (Madrid et al 2018). There are clearly interesting parallels with the situation in marmosets, raising the intriguing possibility that similar selective forces have led to the evolution and maintenance of *SLC6A4* polymorphisms in unrelated primate lineages on different continents.

## Acknowledgements

We thank Nancy Caine, Andrew Kitchener, Jim Dietz, Andrew Smith, Hannah Buchanan-Smith, Gustl Anzenberger, Michael Bruford, Oliver Ryder and Anna Feistner for samples, and Nancy Caine for helpful comments on the manuscript. We thank the University of Cambridge Career Support Scheme for staff funding (HT) and the British Council for funding the trip to Brazil with the Researcher Links travel grant awarded to AMS, in collaboration with Dr Jorge Zeredo at the University of Brasilia, Brazil. We also thank Murray Edwards College, Cambridge, for funding.

## Supplementary Information

Suppl Table 1 Samples

Suppl Table 2 Primers

Suppl Table 3 Summary of other sequence variants (SNPs and indels) detected

