## Supplementary tables for "Evolution of threat response-related polymorphisms at the *SLC6A4* locus in callitrichid primates"

**Supplementary Table 1. Samples examined**

**Species	(Sample size)	LabID	Origin	Source**

Callithrix jacchus (2)	Cja13	National Museums of Scotland	Andrew Kitchener

Cja292	CRES	Oliver Ryder

Callithrix penicillata (7)	Cpe3, Cpe4, Cpe5, Cpe6, Cpe8, Cpe9, Cpe11	Primate Centre, University of	This study

Brasília.

Callithrix geoffroyi (9)	Cge32	National Museums of Scotland	Andrew Kitchener

Geoff, Snap, June, May, April, Cra, Sophia, Shiksa	San Diego Wild Animal Park	Nancy Caine

Mico argentatus (1)	Car5	National Museums of Scotland	Andrew Kitchener

Cebuella pygmaea (3)	Cep592, Cep690	CRES	Oliver Ryder

Cpy3	Zoological Society of London	Michael Bruford

Callimico goeldii (8)	Cgo17	National Museums of Scotland	Andrew Kitchener

Cgo20	Zoological Society of London	Michael Bruford

Cgo1, Cgo2, Cgo3, Cgo4, Cgo2186, Cgo1136	University of Zurich	Gustl Anzenberger

Leontopithecus rosalia	(7)	Lro11	National Museums of Scotland	Andrew Kitchener

Lr413, Lr225, Lr194, Lr653, Lr529, Lr490	Poço das Antas, Brazil	Jim Dietz

Leontopithecus chrysomelas (2)	Lch1, Lch2	Zoological Society of London	Michael Bruford

Leontopithecus chrysopygus (3)	Lcp1, Lcp2, Lcp3	Jersey Zoo	Anna Feistner

Leontocebus fuscicollis (13)	Sfi1, Sfi2, Sfw1, Sfw2, Sfw5, Sfw7	National Museums of Scotland	Andrew Kitchener

SF2214, SF2365, SF296, SF2583, SF2584, SF1497, SF1045	Belfast Zoo	Hannah Buchanan-Smith

Oedipomidas geoffroyi (1)	Sge1	National Museums of Scotland	Andrew Kitchener

Saguinus midas	(2)	Smi2, Smi3	National Museums of Scotland	Andrew Kitchener

Saguinus bicolor (2)	Sbi1, Sbi2	National Museums of Scotland	Andrew Kitchener

Tamarinus labiatus (6)	Sla2, Sla3	National Museums of Scotland	Andrew Kitchener

SL1708, SL656, SL2306, SL1F7	Belfast Zoo	Hannah Buchanan-Smith

**Supplementary Table 2. Primers (All 5’-3’)**

PCR primers

RPRF	Forward	CAGACAACCGTGTTCATCTG	Santangelo et al. 2016

RFRR	Reverse	GATTCTAGTGCCACCTAGAC	Santangelo et al. 2016

RPRFL	Forward	AGCAGACAACCGTGTTCATCTG	This study

RPRRL	Reverse	GATTCTAGTGCCACCTAGACGC	This study

PTF	Forward	CTGCCCCCAGAATAAAATTCC	This study

PTR	Reverse	AGCGGGAGGAGTCAGCAC	This study

Internal sequencing primers

SeqF1	Forward	AGCAGCACCTAACCCTCCTA	Santangelo et al. 2016

SeqF2	Forward	TCCCCACTAGGCATTGCTAC	Santangelo et al. 2016

SeqR1	Reverse	ACTGGCAGGTGGATGTTGAG	This study

SeqLF	Forward	CCCTCCAGCATTCCCTTTGT	This study

**Supplementary Table 3. Intraspecific variants found in the repeat region (excluding the 4 functional SNP sites in *C. jacchus*)**

**Species	(Sample size)	Variants**

Callithrix jacchus (2)	None

Callithrix penicillata (7)	6 SNPs

Callithrix geoffroyi (9)	2 SNPs, one indel (1bp)

Mico argentatus (1)	None

Cebuella pygmaea (3)	4 SNPs, one indel (22bp, one repeat unit)

Callimico goeldii (8)	7 SNPs, one indel (11bp, half of a repeat unit)

Leontopithecus rosalia	(7)	2 SNPs

Leontopithecus chrysomelas (2)	1 SNP

Leontopithecus chrysopygus (3)	3 SNPs

Leontocebus fuscicollis (13)	5 SNPs, one indel (22bp, one repeat unit)

Oedipomidas geoffroyi (1)	None

Saguinus midas	(2)	None

Saguinus bicolor (2)	None

Tamarinus labiatus (6)	2 SNPs
